# acmgscaler: An R package and Colab for standardised gene-level variant effect score calibration within the ACMG/AMP framework

**DOI:** 10.1101/2025.05.16.654507

**Authors:** Mihaly Badonyi, Joseph A Marsh

## Abstract

**Summary:** A genome-wide variant effect calibration method was recently developed under the guidelines of the American College of Medical Genetics and Genomics and the Association for Molecular Pathology (ACMG/AMP), following ClinGen recommendations for variant classification. While genome-wide approaches offer clinical utility, emerging evidence highlights the need for gene- and context-specific calibration to improve accuracy. Building on previous work, we have developed an algorithm tailored to converting functional scores from both multiplexed assays of variant effects (MAVEs) and computational variant effect predictors (VEPs) into ACMG/AMP evidence strengths. Our method is designed to deliver consistent performance across different genes and score distributions, with all variables adaptively determined from the input data, preventing selective adjustments or overfitting that could inflate evidence strengths beyond empirical support. To facilitate adoption, we introduce `acmgscaler`, a lightweight R package and a plug-and-play Google Colab notebook for the calibration of custom datasets. This algorithmic framework bridges the gap between MAVEs/VEPs and clinically actionable variant classification.

**Availability:** The package and the Colab notebook are available on GitHub: https://github.com/badonyi/acmgscaler.

**Supplementary data:** Supplementary data can be downloaded from the OSF repository https://osf.io/7hjnm.

## Background

To establish a unified approach to clinical genetic and genomic testing, the American College of Medical Genetics and Genomics and the Association for Molecular Pathology (ACMG/AMP) outlined guidelines for assessing the pathogenicity and benignity of variants in Mendelian disease genes^1^. These guidelines have since been formalised within a probabilistic framework, demonstrating how ACMG/AMP criteria for combining clinical evidence align with likelihood ratios for pathogenicity^2^. Within this framework, the strength of evidence for each classification (supporting, moderate, strong, and very strong) has been defined to increase exponentially. This model has provided the foundation for systematically validating and calibrating pathogenicity evidence, advancing efforts to refine variant classification and extend the role of predictive tools beyond supporting-level evidence^3^.

One such extension applies to computational variant effect predictors (VEPs), where appropriate weighting of evidence is integral to ensuring accurate classification. Recently, Pejaver et al. developed a method to estimate the pathogenic-over-benign density ratio for a variant by applying a narrow sliding window to a sorted list of scores labelled as pathogenic or benign^4^. The resulting odds of pathogenicity, equivalent to positive likelihood ratios (LR^+^; hereafter referred to as LR), can be compared to evidence thresholds established by the ACMG/AMP framework under a predefined prior probability of pathogenicity^1–3^. Using this approach, the authors estimated genome-wide thresholds, *i*.*e*., score intervals, for each evidence level across thirteen widely used VEPs. This work has since been extended with the calibration of additional VEPs_5_.

Applying genome-wide calibration to VEP scores for variant classification has demonstrated notable benefits. For example, one study used genome-wide thresholds in 300 probands with suspected rare disease and found a 2.6-fold increase in variants reaching strong evidence for pathogenicity compared to when only established disease-associated genes were considered^6^. Importantly, this analysis concluded that genome-wide calibration is unlikely to result in the inappropriate classification of an excessive number of variants as pathogenic or likely pathogenic^6^. However, others have raised concerns about the limitations of genome-wide calibration, particularly in terms of obscuring performance across different genes and modes of inheritance^7–9^. These findings suggest that genome-wide calibration alone may not be sufficient to address these discrepancies and highlight the need for a gene- and context-specific calibration method.

Multiplex assays of variant effect (MAVEs) provide an unbiased source of functional evidence^10–12^, offering huge potential to improve the classification of germline genomic variants, particularly rare missense variants frequently classified as uncertain significance^13^. However, similar to the output of VEPs, raw scores from MAVEs vary in range and distribution across genes and do not readily correspond to ACMG/AMP evidence strengths.

Here, building on previous work^8,14,15^, we develop a method based on bootstrapped kernel density estimates to compute LRs for any functional score distribution—whether derived from MAVEs or VEPs—using pathogenic and benign truthset labels. The resulting LRs support robust and reproducible integration of functional scores from diverse sources into variant classification under ACMG/AMP guidelines. We have made our method available as an R package for high-throughput calibration studies, alongside a Google Colab notebook providing a simple interface for bench scientists.

## Implementation

### Likelihood ratio estimation

We follow van Loggerenberg et al. in using probability density estimation with a Gaussian smoothing kernel to derive LRs for functional scores^14^, as the sliding window method developed by Pejaver et al.^4^ is not feasible at the gene level due to data sparsity. We set a lower limit on the number of variants in the truthset, requiring at least ten benign and ten pathogenic labels each. To avoid extrapolating LRs beyond the observed range, input scores are first clamped to the range of labelled scores, and then rescaled to the [0, 1] interval to ensure scale invariance. Density estimation is performed separately on the pathogenic and benign score distributions, using 1,024 equidistant evaluation points and a bandwidth selected by biased cross-validation. This process is repeated 1,000 times by resampling the reference distributions with replacement, applying kernel density estimation to each resample and projecting the resulting densities onto a common grid of 1,024 points via linear interpolation. A fixed random seed is used to eliminate run-to-run variability.

Log-LRs are computed across resamples as the difference between pathogenic and benign log-densities at each index in the common grid, resulting in a 1,000 × 1,024 matrix. To stabilise estimates in sparse regions, we apply adaptive regularisation:

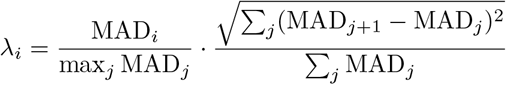

Here, MAD_*i*_ is the median absolute deviation of log-LRs at grid index *i*. The first term scales MAD values to reflect relative variability across the score range. The second term, a root-sum-of-squares of lagged differences, acts as a roughness penalty similar to L2 regularisation. This formulation penalises sharp or erratic fluctuations in the log-LRs while leaving stable regions largely unaffected.

At this stage, the log-LRs may be non-monotonic due to boundary effects or multimodal structures in the score distributions, which could reflect noise, assay or VEP limitations, or inaccuracies in truthset labels. When higher scores indicate a greater likelihood of pathogenicity, a lower score should not yield a higher LR, as this would suggest local overfitting. To correct this, we apply isotonic regression to enforce monotonicity in the log-LR matrix, ensuring a gradual or stepwise increase in evidence across the score range. The direction of monotonicity is determined by the sign of the Spearman correlation between the column-wise mean of log-LRs and the input scores. The final LRs are obtained by computing the median log-LR across resamples, exponentiating to return to linear space, and mapping the estimates to the input scores based on their grid indices. A flowchart illustrating the method is shown in **Figure 1**.

**Figure 1.**
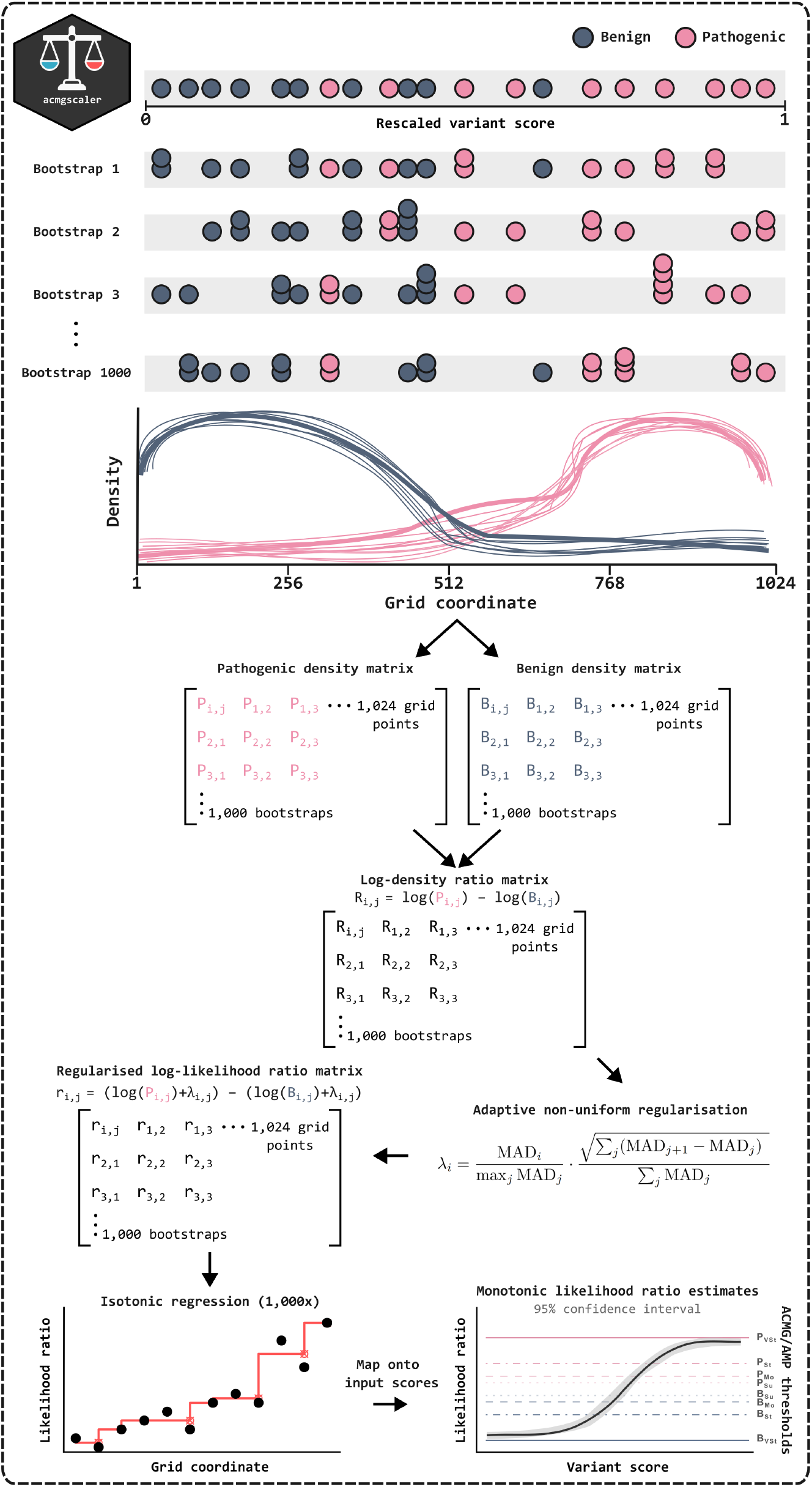
Method flowchart.

### Computing confidence intervals for the likelihood ratios

We compute 95% confidence bounds for the population median of log-LRs across resamples by inverting the sign test, using the exact binomial distribution of order statistics^16^. An intuitive definition of this confidence interval is that if calibration were repeated 100 times with different random seeds, the median LR would fall within the interval in 95 of those repeats. The choice of a confidence interval for the location of the median log-LR, rather than, *e*.*g*., a percentile interval for the distribution of log-LRs, better aligns with our classification goal, as we care most about where the typical (central) log-LR lies. It is important to clarify that these confidence bounds do not represent uncertainty in the posterior probability of pathogenicity as defined in the ACMG/AMP framework. Instead, they reflect variability in the LR itself, estimated from the regularised and monotonic log-LR distribution. A classification based on a variant for which both the point estimate and the lower (for pathogenic) or upper (for benign) confidence bound exceed a given threshold is less likely to be affected by future truthset updates, *e*.*g*., the inclusion of additional pathogenic variants, than one based on the point estimate alone. Accordingly, we classify variants into evidence strengths using the lower bound when the point estimate exceeds one, and the upper bound when it falls below.

### Minimum truthset size requirement

While as few as four samples can keep relative error low in one-dimensional kernel density estimation^17^, we require a minimum truthset of ten pathogenic and ten benign variants to ensure broad generalisability across diverse distributions. We evaluated this choice using Monte Carlo cross-validation. Specifically, we selected genes (N=187) with at least twenty missense variants classified as benign/likely benign (B/LB) and pathogenic/likely pathogenic (P/LP) in ClinVar^18^, requiring a minimum review status of one star. From each gene, we randomly sampled ten variants per class without replacement and calibrated AlphaMissense^19^ and CPT-1^20^ scores, which were selected based on their top performance in a recent benchmarking study^21^. We repeated this process for 2,000 trials. For each trial, we calculated the observed LR as the ratio of true pathogenic to true benign variants in a given ACMG/AMP evidence level, excluding those used for calibration. We then divided this value by the background ratio of true pathogenic to true benign variants in the gene.

The aim of this analysis is to assess how well the calibrated thresholds recover expected evidence strengths when using the minimum sample size. For benign evidence, observed LRs should typically be at or below the expected value, and for pathogenic evidence, at or above it. This would show that the minimum sample size is not prone to inflating evidence strength due to chance variation in small truthsets. In **Figure 2A**, we compare the observed LRs to the expected LR under a prior probability of 0.1. For the vast majority of genes, the observed LRs follow this pattern across the range from strong benign to strong pathogenic evidence. Caution is warranted for the very strong category, which is harder to assess due to data sparsity and occasional instability in the estimates. Nevertheless, these results demonstrate that the calibrated thresholds are generally reliable with the default minimum truthset size requirement.

**Figure 2.**
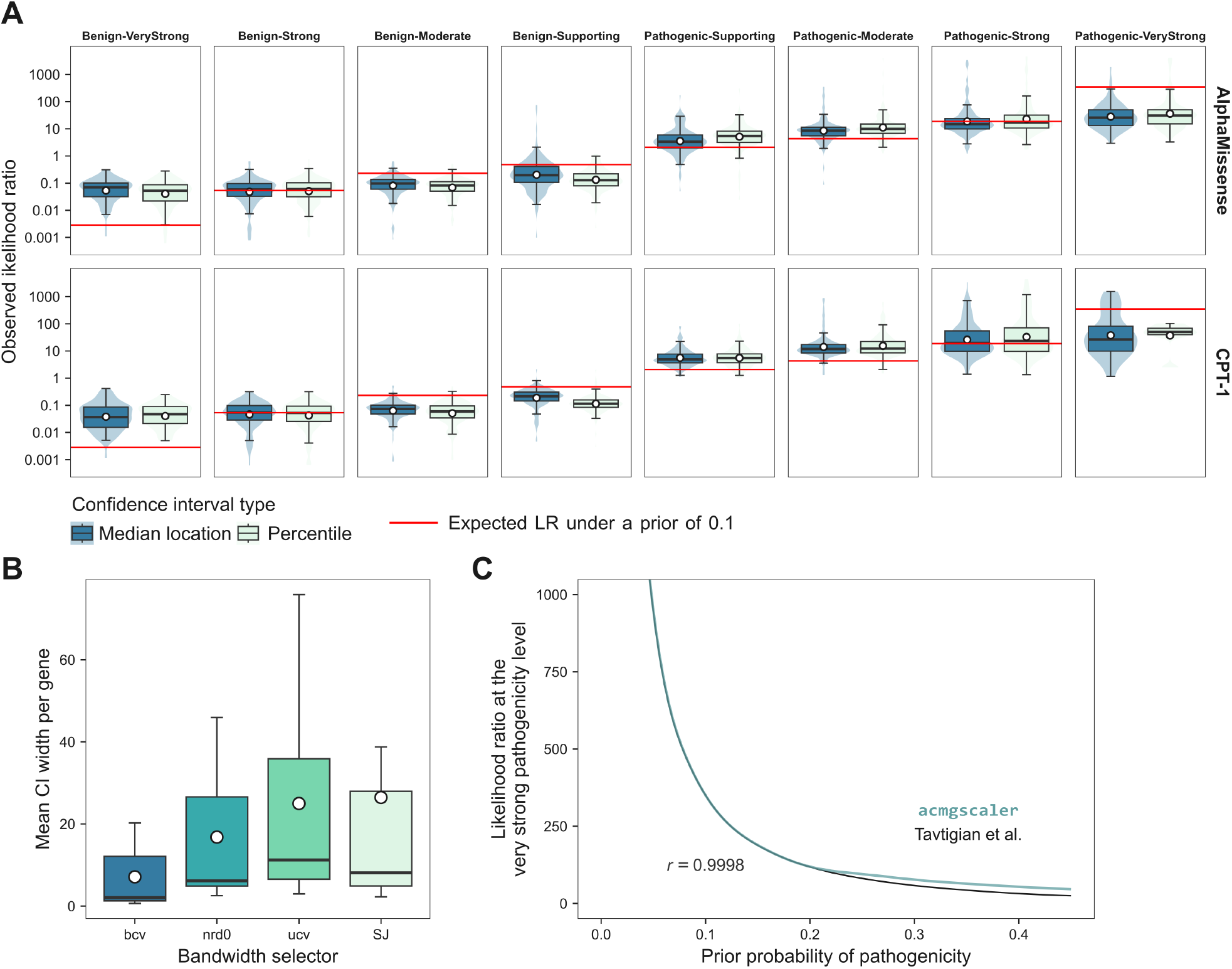
**A**: Box-violin plots of the Monte Carlo cross-validation showing the distribution of observed LRs for 187 genes across ACMG/AMP evidence strengths and for AlphaMissense and CPT-1 scores separately. Each evidence strength category shows observed LRs with the classification either based on the median location or the percentile method for calculating confidence intervals. Boxes represent data within the 25th and 75th percentiles, with the middle line indicating the median, dots indicating the mean, and whiskers extending to 1.5× the interquartile range. **B**: Comparison of bandwidth selection methods for computing LRs. The *y* axis shows the gene-averaged width of the LR confidence interval across calibrated MAVE, AlphaMissense and CPT-1 scores. **C**: Likelihood points for the very strong pathogenicity level as implemented in the `acmgscaler` package. The Pearson correlation (*r*) with the reference method by Tavtigian et al. is shown.

Additionally, we used this simulation framework to compare the median location-based and percentile methods for computing confidence intervals. As shown in **Figure 2A**, the resulting distributions differ only marginally, supporting the adoption of the median location-based method.

### Choice of bandwidth selector

The bandwidth parameter controls the smoothness of the estimated density functions and can substantially affect the resulting LRs. We evaluated four automatic bandwidth selectors available in base R: ‘nrd0’ (Silverman’s rule of thumb), ‘bcv’ (biased cross-validation), ‘SJ’ (Sheather–Jones), and ‘ucv’ (unbiased cross-validation)^17,22,23^. To select the most suitable method, we compared the average width of the LR confidence interval across truthset variants, which directly contribute to the density estimates, for seven human disease genes with available MAVE, AlphaMissense, and CPT-1 scores. All seven genes include at least ten missense variants classified as B/LB and P/LP pathogenic in ClinVar, with a minimum review status of one star, which served as truthset labels for calibration. For each gene, the MAVE scoreset with the highest area under the receiver operating characteristics curve was selected. The genes, MAVE studies, scoreset sources, and the number of pathogenic and benign variants are listed in **Table S1**. As shown in **Figure 2B**, ‘bcv’ yielded the narrowest confidence intervals on average, suggesting greater stability, and was therefore set as the bandwidth selector in our method.

### Determining likelihood points for a given prior

The prior probability of pathogenicity (hereafter ‘prior’) represents the estimated likelihood that a randomly selected, previously uncharacterised variant is disease-causing in a patient presented with a given phenotype before considering variant-specific evidence. Global estimates, such as 0.1^2^and the more recent 0.0441^4^, provide a useful baseline across diverse disease-associated genes, but priors may vary substantially between genes, and even among phenotypes. This variation reflects differences in gene properties, including their biological role, degree of functional constraint, associated disease prevalence, mode of inheritance, and molecular mechanism. To leave room for future advances in gene-specific prior estimation, the default prior of 0.1 can be replaced with a user-specified value.

Likelihood points corresponding to ACMG/AMP evidence strengths under a predefined prior can be determined using the framework introduced by Tavtigian et al^2^. Here, we approximate this by considering that the *P*_*ii*_ and *LP*_*i*_ combining rules yield internally inconsistent likelihood points: *P*_*ii*_ produces a posterior probability below the expected 0.99 threshold, whereas *LP*_*i*_ exceeds it. To determine the very strong pathogenicity threshold, we identify the smallest LR that satisfies at least 13 of 14 posterior probability criteria across the *LP* and *P* combining rules. Although the original ACMG/AMP framework imposes additional conditions—requiring the *LB* combining rules to have a posterior probability between 0.05 and 0.001, and the *B* combining rule to be below 0.001—we find that enforcing the criteria for *P* and *LP* implicitly maintains a similar relationship between the prior and the LR threshold for the very strong pathogenicity evidence (P_*VSt*_). As shown in **Figure 2C**, for priors below 0.2, our method closely aligns with the proposed combining rules, yielding slightly more conservative values above 0.2. Notably, for the prior of 0.1 used by Tavtigian et al., which gives a P_*VSt*_ threshold of 350, our approach yields the same value. For the prior of 0.0441 suggested by Pejaver et al., which corresponds to a P_*VSt*_ threshold of 1124, our estimate is 1131. A difference of seven LR points is negligible in practice, as it represents only 0.64% of the range between the strong and very strong evidence thresholds under this prior.

### Deriving score thresholds for evidence levels

To compute score intervals for the evidence levels, we first determine P_*VSt*_ under the specified prior and derive the corresponding likelihood points for the remaining levels. For each evidence strength, the score threshold is obtained by linearly interpolating the score at which the lower confidence bound of the LR crosses the evidence threshold if the point estimate exceeds one, and the upper bound otherwise. Importantly, the interpolation is done on the common grid coordinates, ensuring that the values are derived from the full resolution of the LR curve.

### Calibration examples

Full calibration profiles for the seven genes, using a global prior of 0.1, are shown separately for MAVE assays, AlphaMissense, and CPT-1 scores in **Figure 3**. Despite substantial differences in distribution shape, modality, and scale across score sets, the LR estimates generally track well with the empirical density distributions of pathogenic and benign variants. This alignment suggests that the method is robust to underlying functional score characteristics and captures consistent evidence strengths across both experimental and computational predictors. While the confidence intervals may appear narrow, this reflects the robustness of the central tendency of the LR estimates. As a complementary visualisation, we show percentile-based confidence intervals in **Figure S1**, which better reflect the overall variance of the resampled log-LR distribution but are less aligned with our classification objective, which depends on the location and uncertainty of the central estimate. Calibrated datasets are available on OSF (https://osf.io/7hjnm).

**Figure 3.**
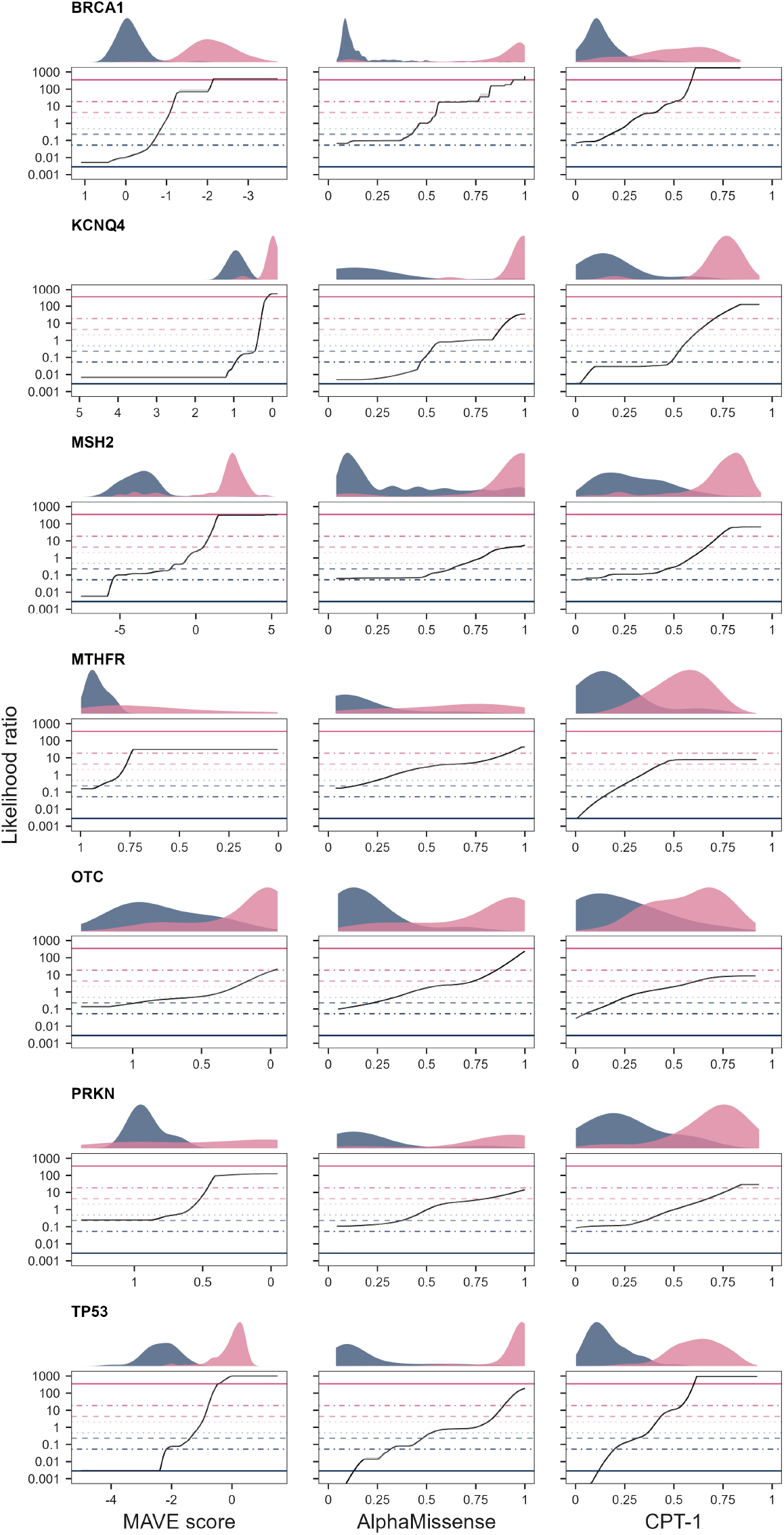
Calibrated MAVE assay, AlphaMissense and CPT-1 scores for missense variants in human genes using `acmgscaler`. Density plots show the distribution of pathogenic (pink) and benign (blue) scores. In MAVE assays, the *x* axis is inverted for BRCA1, KCNQ4, MTHFR, OTC, and PRKN for consistent directionality. Where visible, shaded areas represent 95% confidence intervals. Horizontal bars show evidence strengths under a prior of 0.1, ranging from pathogenic very strong at the top to benign very strong at the bottom. Note that the *x* axis is log-scaled with raw LR tick labels, such that log-LR = 0 corresponds to LR = 1 (where the odds of pathogenicity equal those of benignity, defining the log-midpoint of the Indeterminate evidence level).

To showcase context-specific calibration, we examined the MTHFR assay conducted in a p.Ala222Val background at different folinate concentrations^24^. The p.Ala222Val variant is associated with reduced enzyme activity and thermostability^25^, yet it is also common, with carrier frequencies approaching 50% in some ancestry groups. Individuals homozygous for the allele display elevated homocysteine levels (hyperhomocysteinemia), a hallmark of folate-cycle disruption, which can be ameliorated by folate supplementation^26^. The p.Ala222Val background captures both genetic and nutritional context-dependent effects, which are essential for accurate functional annotation and clinical interpretation of other MTHFR missense variants. To calibrate these assays, we used early-onset hyperhomocysteinaemia variants collected by the authors, and appended ClinVar benign variants with population variants observed at least once in the homozygous state in gnomAD^27^. This truthset makes it possible to run the calibration separately on the catalytic and regulatory domains, with both meeting the minimum truthset size requirement.

**Figure 4A-B** shows the result of the calibration. Consistent with previous findings, scores in the regulatory domain tend to be predictive only of pathogenicity, whereas scores in the catalytic domain can support moderate evidence of pathogenicity and strong evidence of benignity. While the original study reported stronger pathogenic evidence from catalytic domain scores, this discrepancy may reflect differences in the truthset or the effect of our more conservative regularisation, which penalises high-variance estimates more strongly. Despite this, we observe a clear folinate concentration-dependent effect on ACMG/AMP evidence strengths: lower concentrations yield a higher fraction of missense variants classified as pathogenic and a lower fraction classified as benign, both with a shift towards stronger levels of evidence (**Figure 4C**). Thus, our method successfully recapitulates gene-specific effects observed in the MTHFR study, and illustrates how incorporating additional context into calibration frameworks can improve variant interpretation.

**Figure 4.**
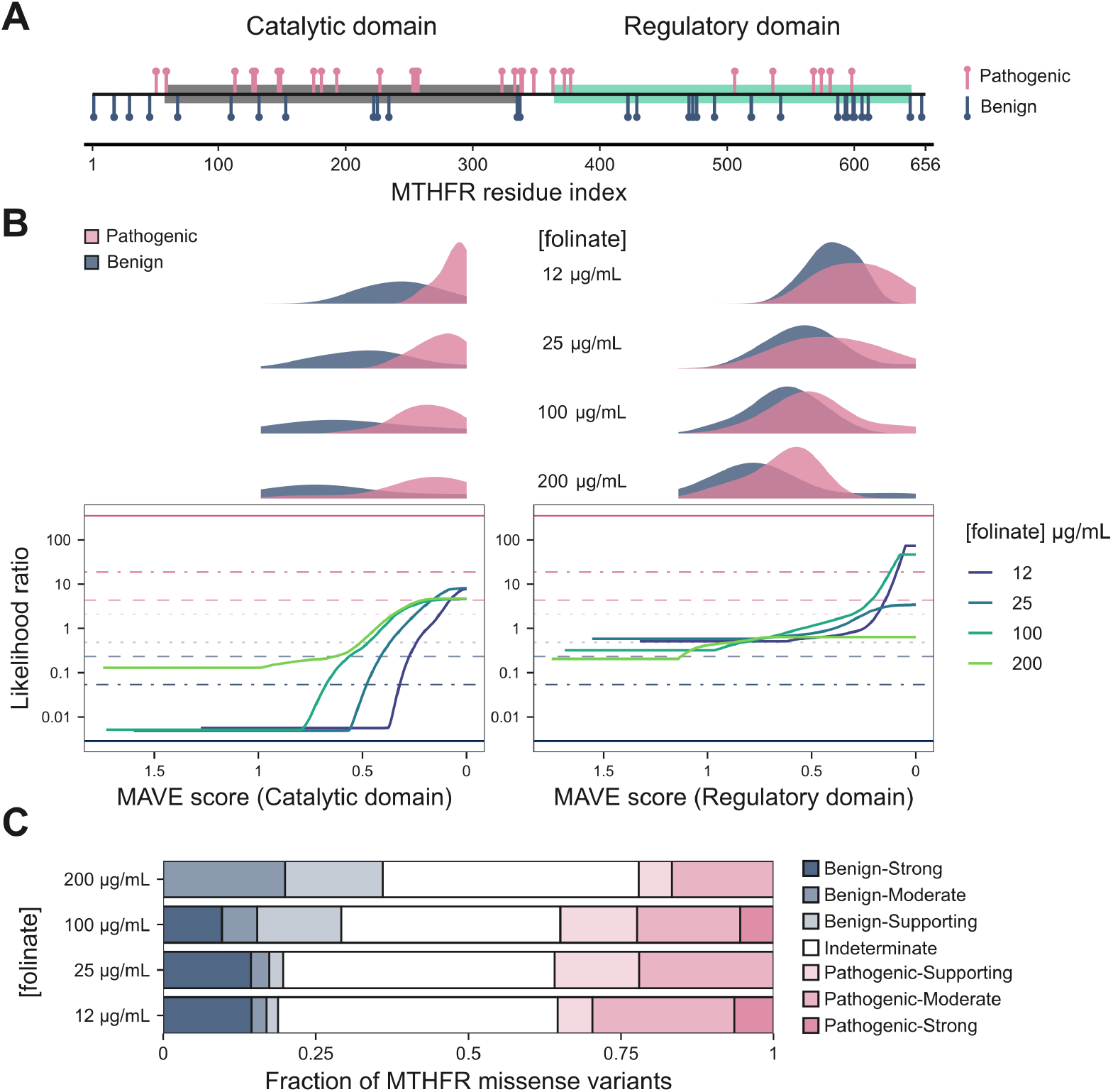
Calibration results for MTHFR MAVE scores in the p.Ala222Val background conducted at different folinate concentrations by Weile et al. (2021). **A**: Location of pathogenic and benign truthset variants along the MTHFR sequence, with the catalytic and regulatory domains highlighted. **B**: MAVE score *vs* LR plots at different folinate concentrations. The density distributions of truthset variants are shown on the top. **C**: Composition of ACMG/AMP evidence classes for all possible MTHFR missense variants with a MAVE score, subset by folinate concentration.

### Genome-wide *vs* gene-specific thresholds

In **Figure 5** we compare genome-wide AlphaMissense thresholds^5^ with gene-specific thresholds derived for the seven genes shown in **Figure 3**. Notably, gene-specific calibration reveals that MSH2 and PRKN do not attain strong pathogenicity evidence, and that BRCA1, MSH2, MTHFR, OTC and PRKN likewise fail to reach strong benign evidence. In contrast, BRCA1 reaches very strong pathogenic evidence and TP53 reaches very strong benign evidence when thresholds are determined at the gene level. Overall, genome-wide and gene-specific thresholds vary substantially: in every case where both exist, the genome-wide benign threshold is lower. Moreover, in KCNQ4, MSH2, and TP53 the supporting pathogenicity thresholds are slightly higher than their genome-wide equivalents. This pattern suggests that, while genome-wide thresholds are unlikely to overcall pathogenic variants in these genes, adopting gene-specific thresholds would enable a greater proportion of variants to be classified as benign. These results further stress the importance of considering the unique functional landscapes of genes when applying calibration thresholds.

**Figure 5.**
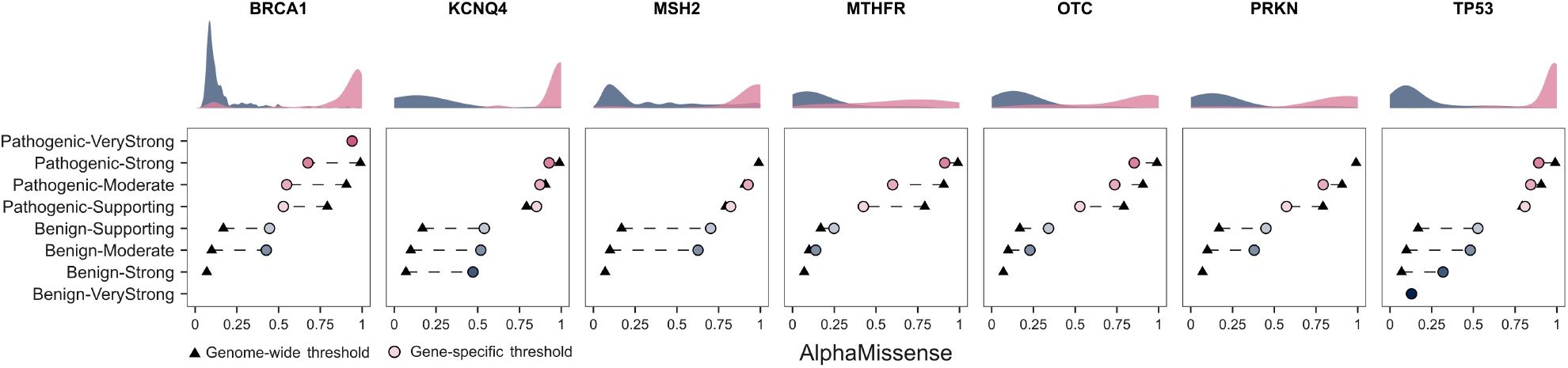
Genome-wide *vs* gene-specific AlphaMissense thresholds for the seven genes shown in **Figure 3.** Density plots show the distribution of pathogenic (pink) and benign (blue) scores. Dashed lines represent a shift between genome-wide and gene-specific thresholds.

## Colab notebook

We have created a Google Colab interface for `acmgscaler`, allowing calibration of custom datasets without the need to install R. The input form, along with an example calibration output, is shown in **Figure 6**. The available settings are simplified to a numeric score column and a class column containing the truthset labels. The class column should include at least ten instances of ‘P’ (pathogenic/positive class) and ‘B’ (benign/negative class) labels each. For any other or missing labels, LRs will also be computed. A custom prior can be set, influencing the evidence strength-based classification results. Upon running the cell, an upload button appears to provide data in CSV format. Once calibration is complete, the Colab automatically downloads the results, while the interactive table allows data export in different formats or filtering through a simple graphical interface. A separate cell enables visualisation and export of the input score *vs* LR curve, with the corresponding ACMG/AMP evidence thresholds highlighted under the predefined prior. The score thresholds can be displayed and exported for downstream use.

**Figure 6.**
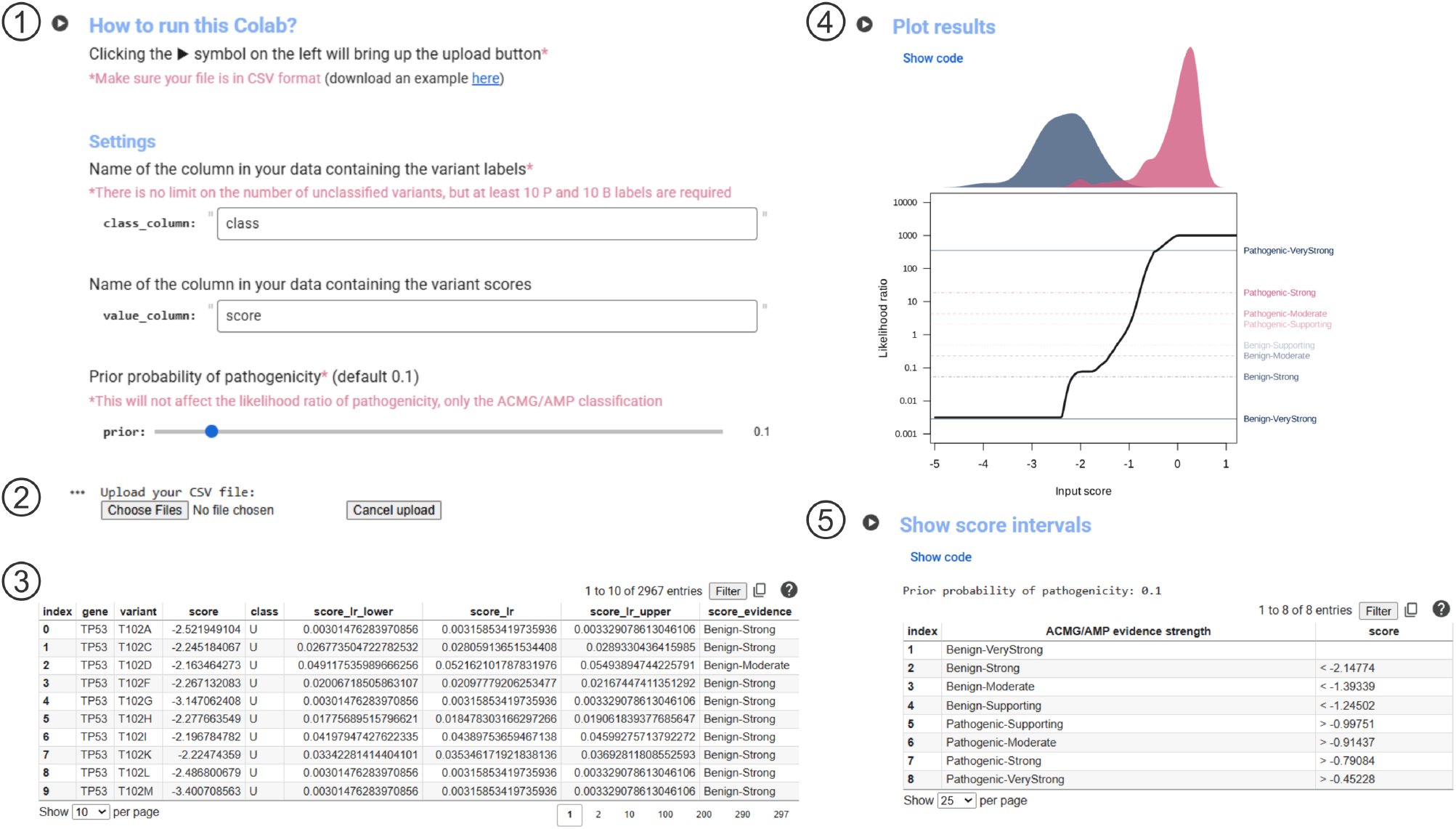
Colab notebook for functional score calibration and variant classification. **1)** Input fields. **2)** Upload button. **3)** Interactive results table. **4)** Visualisation of the input score *vs* LR curve. **5)** Display of score thresholds for evidence levels.

## R package

### Installation

The development version of `acmgscaler` can be installed using `devtools`:

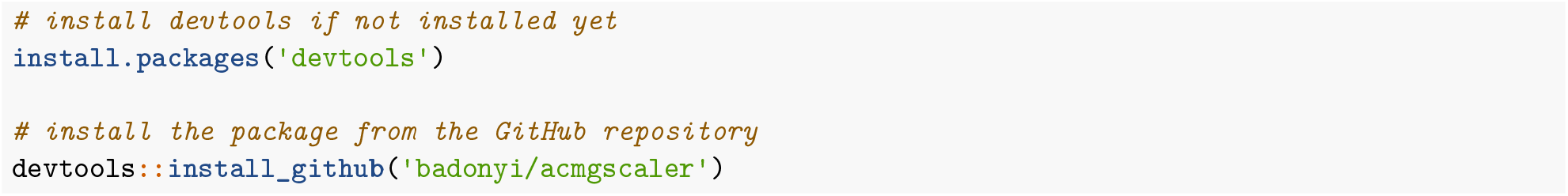

### Usage

The main function `calibrate()` adds LRs as new columns to an existing dataframe containing the variant scores and the truthset labels, and derives the score thresholds for the ACMG/AMP evidence strengths. For convenience, these are computed for all numeric columns if `value` is set to NULL. The function returns a named list with two elements:

- `likelihood_ratios`: the input dataframe augmented with LR estimates and their confidence bounds (`<value>_lr_lower, <value>_lr, <value>_lr_upper`), and a column with assigned ACMG/AMP classifications under the set prior (`<value>_evidence`).
- `score_thresholds`: the lower and upper score bounds defining each evidence strength.

If a grouping variable is supplied, `calibrate()` returns a nested list of the same structure for each group. The example data included in the package contains MAVE scores for BRCA1 and TP53 variants and their class labels from this study.

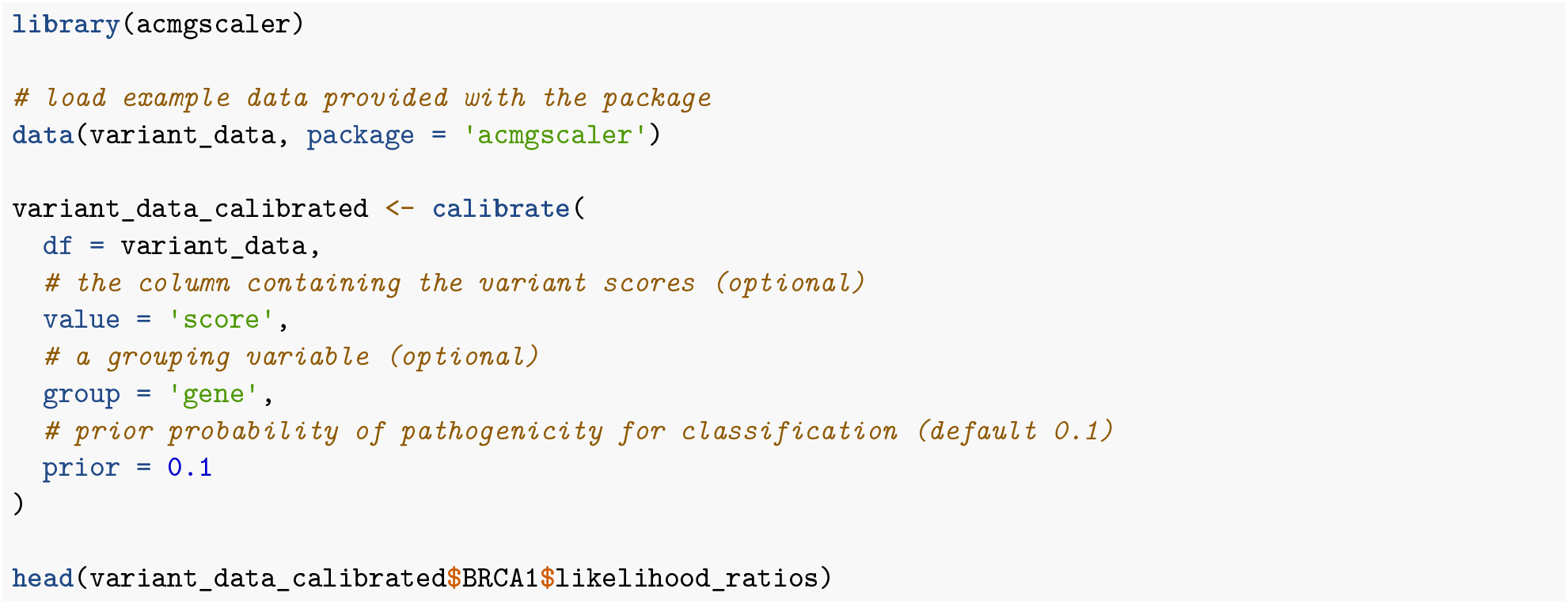

| gene | variant | class | score | score_lr_lower | score_lr | score_lr_upper | score_evidence |
| --- | --- | --- | --- | --- | --- | --- | --- |
| BRCA1 | L3F | B | -0.4736 | 0.0230 | 0.0241 | 0.0256 | Benign-Strong |
| BRCA1 | R7L | B | -0.0165 | 0.0079 | 0.0092 | 0.0103 | Benign-Strong |
| BRCA1 | V11I | B | -0.3134 | 0.0156 | 0.0165 | 0.0174 | Benign-Strong |
| BRCA1 | N13K | B | -0.1705 | 0.0129 | 0.0133 | 0.0136 | Benign-Strong |
| BRCA1 | M18K | P | -2.3905 | 397.6330 | 422.8675 | 451.8950 | Pathogenic-VeryStrong |
| BRCA1 | M18L | B | -0.0593 | 0.0083 | 0.0105 | 0.0112 | Benign-Strong |

## Discussion

We developed a functional score calibration method within the ACMG/AMP framework. This approach is entirely data-driven, eliminating arbitrary thresholds and manual tuning, and yields stable LRs across diverse assay types and computational predictors, regardless of distribution shape or scale.

Our methodology builds on the core idea behind `maveLLR`, a kernel density estimation method developed to assign ACMG/AMP evidence strengths for MAVE assays^14,15^. We extend this framework with resampling, which permits the use of a variance-based penalty to shrink unstable estimates towards one and to compute confidence bounds. Monotonicity constraints on the LRs are enforced via isotonic regression, introducing only modest, local adjustments to the curve and preserving its empirically supported structure. The combination of adaptive shrinkage and a simplified thresholding procedure ensures that the assigned evidence strengths remain robust and conservative. These steps are implemented entirely in base R using highly optimised core functions, maintaining long-term backwards and forwards compatibility. Moreover, we provide a Google Colab wrapper, bypassing the need to install R for single custom datasets.

Zeiberg et al. recently introduced an alternative calibration framework for MAVE assays based on a skew-normal mixture model^28^. This method fits score distributions using parametric components, with monotonicity enforced via an expectation-maximisation algorithm. By jointly estimating parameters across the entire score distribution, this approach is particularly effective at stabilising sparse or extreme score regions. The authors also propose a strategy for estimating the prior (the probability that a variant in a patient with a given phenotype is causal for the disease, before incorporating evidence) by modelling gnomAD^27^ scores as a mixture of pathogenic and benign components. In contrast, our framework computes LRs directly from empirical distributions of labelled variants, without relying on parametric assumptions, making it readily generalisable to any continuous score set, including those of VEPs. A standardised calibration pipeline for both MAVE and VEP data ensures all functional scores are treated identically, making evidence integration and auditing straightforward within the ACMG/AMP framework.

The sole parameter in our method is the prior. While genome-wide priors have been estimated before^2,4^, deriving reliable context-specific priors remains challenging. This is because Mendelian disease genes frequently have multiple phenotypes, which may differ in their population prevalence, mutational spectrum^29^ and molecular mechanism^30^. Even in genes associated with a single phenotype, estimating priors from population data, such as gnomAD, can be problematic. For example, genes with recessive conditions typically have a limited number of alleles that have been observed in the homozygous state, which represent a more suitable proxy for benign variants in these genes than heterozygous alleles^31^. We anticipate that with increasing population sequencing data and advances in statistical genetics methods, reliable estimation of context-specific priors suitable for practical application will be feasible in the near future. Until then, we recommend using a constant prior, *e*.*g*. 0.1, to avoid the risk of inflating evidence strengths as a result of overestimating the prior.

We developed our method with missense variants in mind, as most VEPs are limited to them. However, the approach can in principle be applied to any variant type, provided the scores are derived from the same assay or VEP and reflect quantitatively comparable functional consequences. Particular care is needed when including nonsense variants from MAVEs. For the majority of MAVE approaches that rely on exogeneous constructs, a premature stop codon will result in a truncated protein product, which may not mimic the biological effects within cells where nonsense-mediated decay is often involved. Similarly, certain MAVEs may not detect aberrant splicing events^32^; thus users should evaluate whether the assay setup captures the relevant biology for each variant type when performing calibration.

Another important consideration is the potential for circularity in the calibration dataset. We used ClinVar-labelled pathogenic and benign variants as the truthset to calibrate MAVE and VEP scores across seven human genes. However, many of the MAVE assays may have already contributed functional evidence towards these clinical classifications^33,34^, raising the possibility of inflating the calibrated evidence strength. VEPs are similarly susceptible to this form of bias^35^. For example, AlphaMissense was trained in part on population allele frequencies, which inform benignity assessments under ACMG/AMP guidelines^36^, although the resulting bias appears modest^21,37^. CPT-1 was partially trained on MAVE data, including the assay for the MTHFR gene, yet showed no strong evidence of bias in our calibration. To mitigate circularity, truthsets ideally should exclude variants whose classification relied on the predictive features being evaluated. As MAVEs and VEPs are increasingly incorporated into clinical variant interpretation, such circularity is likely to become more pervasive. It will therefore be essential to track the provenance of supporting evidence to ensure robust calibration and reliable downstream use. Sources of potential circularity warrant careful consideration in calibration projects intended to inform clinical decision-making.

For simplicity, we used ClinVar-labelled pathogenic and benign variants without stratifying by disease phenotype or inheritance mode. However, such context-specific considerations are critical for accurate clinical calibration and are already adopted by variant curation expert panels. Researchers with gene-specific knowledge often have access to additional pathogenic labels from alternative databases, detailed literature reports, patient pedigrees, and case-control studies. While author-curated reference sets may suffice for exploratory analyses, calibration to clinical genetics standards requires high-quality disease variants defined through expert consensus. Nonetheless, the availability of bona fide benign labels is often more limiting. While allele frequency cutoff-based approaches may yield useful proxy-benign variants for highly constrained dominant genes^38–40^, caution is required for recessive conditions where heterozygous population variants can drive disease. Although VEP-based thresholds could offer an alternative, in supervised models they depend on the training data, which can thus influence the resulting calibration in ways that overstate confidence for variants similar to the training set while underestimating risk for novel or underrepresented variants. Ultimately, large-scale population sequencing remains the most viable avenue to augment benign reference sets and accelerate the discovery of disease-associated variants.

## Notes on system requirement

The package has no external dependencies and has been tested on R versions >=3.5.0. Memory usage is approximately constant owing to the fixed size of the log-LR matrix. Calibrating the datasets in this study required a peak RAM of 150 MB on an Intel(R) Xeon(R) Gold 6338 CPU @ 2.00GHz with 1 GB of RAM.

## Acknowledgements

The authors thank Yifei Shang and Hasan Çubuk for their valuable feedback. This project was supported by funding from the European Research Council (ERC) under the European Union’s Horizon 2020 research and innovation programme (grant agreement No. 101001169) and by funding from the Medical Research Council (MRC) Human Genetics Unit core grant (MC_UU_00035/9).

---

**Figure S1.**
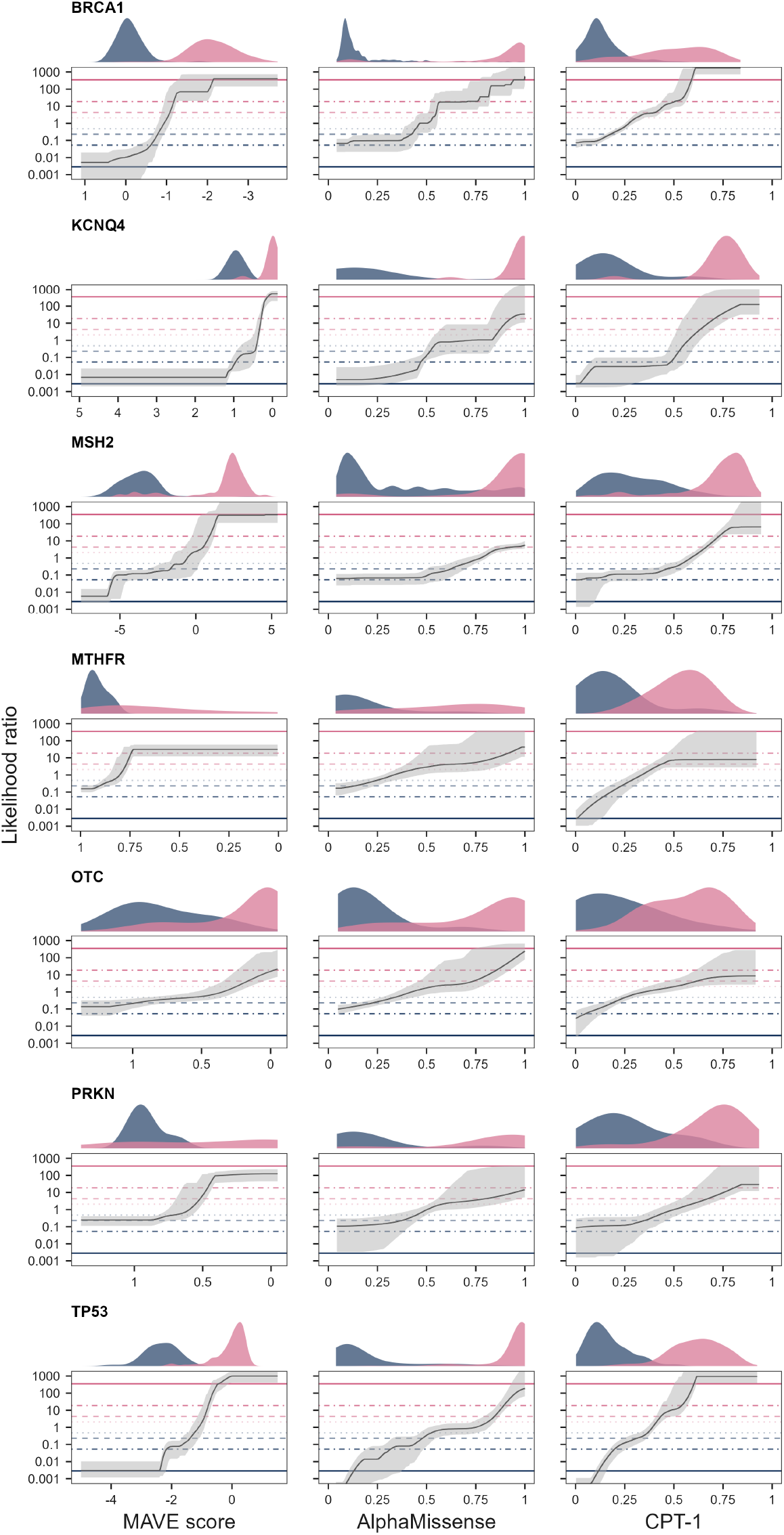
Calibration results as shown in **Figure 3**, with shaded areas representing the 2.5th and 97.5th percentiles of the bootstrap distribution of LR estimates.

**Table S1.**
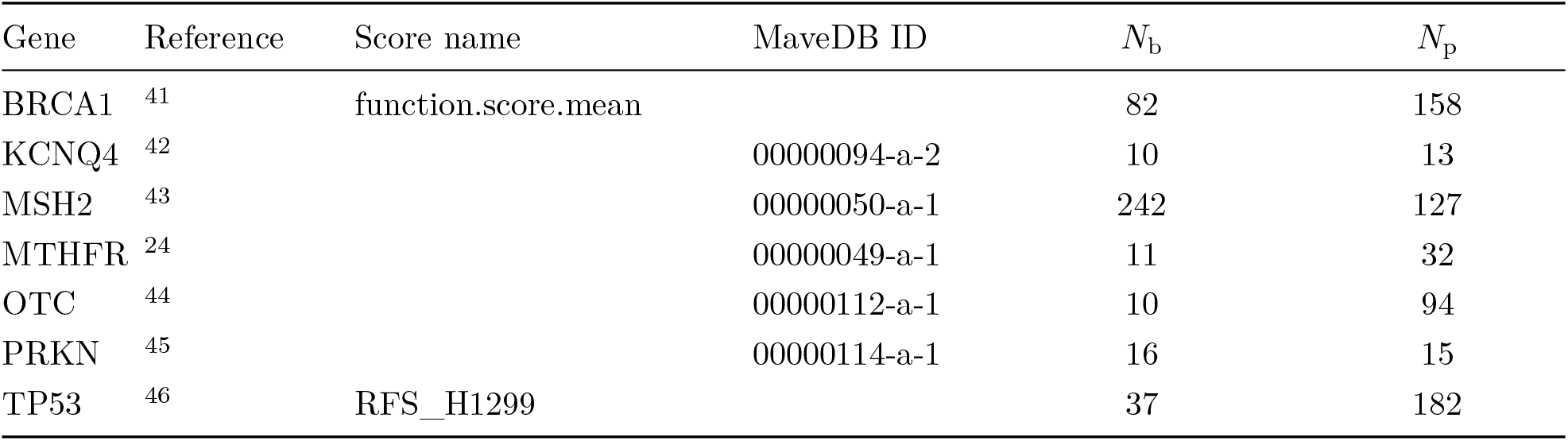
Sources of MAVE datasets calibrated in **Figure 3**. Either the MaveDB ID or the score name from the original publication is given. *N*_b_ and *N*_p_ are the number of benign and pathogenic missense mutations in the datasets, respectively. Note that AlphaMissense and CPT-1 have full coverage in these genes with equal or greater labelled variant counts.

## References

1. Richards, S. et al. Standards and guidelines for the interpretation of sequence variants: A joint consensus recommendation of the american college of medical genetics and genomics and the association for molecular pathology. Genetics in medicine 17, 405–423 (2015).

2. Tavtigian, S. V. et al. Modeling the ACMG/AMP variant classification guidelines as a bayesian classification framework. Genetics in medicine 20, 1054–1060 (2018).

3. Brnich, S. E. et al. Recommendations for application of the functional evidence PS3/BS3 criterion using the ACMG/AMP sequence variant interpretation framework. Genome medicine 12, 1–12 (2020).

4. Pejaver, V. et al. Calibration of computational tools for missense variant pathogenicity classification and ClinGen recommendations for PP3/BP4 criteria. The American Journal of Human Genetics 109, 2163–2177 (2022).

5. Bergquist, T. et al. Calibration of additional computational tools expands ClinGen recommendation options for variant classification with PP3/BP4 criteria. Genetics in Medicine 101402 (2025).

6. Stenton, S. L. et al. Assessment of the evidence yield for the calibrated PP3/BP4 computational recommendations. Genetics in Medicine 26, 101213 (2024).

7. Tejura, M. et al. Calibration of variant effect predictors on genome-wide data masks heterogeneous performance across genes. The American Journal of Human Genetics 111, 2031–2043 (2024).

8. Isakov, O., Fluss, R., Marek-Yagel, D. & Ben Shachar, S. The impact of clinical and molecular variant properties on calibration and performance of variant effect prediction tools. bioRxiv 2024–10 (2024).

9. Dias, M., Orenbuch, R., Marks, D. S. & Frazer, J. Toward trustable use of machine learning models of variant effects in the clinic. The American Journal of Human Genetics 111, 2589–2593 (2024).

10. Weile, J. & Roth, F. P. Multiplexed assays of variant effects contribute to a growing genotype– phenotype atlas. Human genetics 137, 665–678 (2018).

11. Fowler, D. M. et al. An atlas of variant effects to understand the genome at nucleotide resolution. Genome biology 24, 147 (2023).

12. Rubin, A. F. et al. MaveDB 2024: A curated community database with over seven million variant effects from multiplexed functional assays. Genome Biology 26, 13 (2025).

13. Chen, E. et al. Rates and classification of variants of uncertain significance in hereditary disease genetic testing. JAMA network open 6, e2339571–e2339571 (2023).

14. Loggerenberg, W. van et al. Systematically testing human HMBS missense variants to reveal mechanism and pathogenic variation. The American Journal of Human Genetics 110, 1769–1786 (2023).

15. Gebbia, M. et al. A missense variant effect map for the human tumor-suppressor protein CHK2. The American Journal of Human Genetics 111, 2675–2692 (2024).

16. Conover, W. J. Practical Nonparametric Statistics. (John Wiley & Sons, 1999).

17. Silverman, B. W. Density Estimation for Statistics and Data Analysis. (Routledge, 2018).

18. Landrum, M. J. et al. ClinVar: Improvements to accessing data. Nucleic acids research 48, D835– D844 (2020).

19. Cheng, J. et al. Accurate proteome-wide missense variant effect prediction with AlphaMissense. Science 381, eadg7492 (2023).

20. Jagota, M. et al. Cross-protein transfer learning substantially improves disease variant prediction. Genome Biology 24, 182 (2023).

21. Livesey, B. J. & Marsh, J. A. Variant effect predictor correlation with functional assays is reflective of clinical classification performance. Genome Biology 26, 104 (2025).

22. Sheather, S. J. & Jones, M. C. A reliable data-based bandwidth selection method for kernel density estimation. Journal of the Royal Statistical Society: Series B (Methodological) 53, 683–690 (1991).

23. Sain, S. R., Baggerly, K. A. & Scott, D. W. Cross-validation of multivariate densities. Journal of the American Statistical Association 89, 807–817 (1994).

24. Weile, J. et al. Shifting landscapes of human MTHFR missense-variant effects. The American Journal of Human Genetics 108, 1283–1300 (2021).

25. Frosst, P. et al. A candidate genetic risk factor for vascular disease: A common mutation in methylenetetrahydrofolate reductase. Nature genetics 10, 111–113 (1995).

26. Marini, N. J. et al. The prevalence of folate-remedial MTHFR enzyme variants in humans. Proceedings of the National Academy of Sciences 105, 8055–8060 (2008).

27. Chen, S. et al. A genomic mutational constraint map using variation in 76,156 human genomes. Nature 625, 92–100 (2024).

28. Zeiberg, D. et al. Gene-based calibration of high-throughput functional assays for clinical variant classification. bioRxiv 2025–04 (2025).

29. Chong, J. X. et al. The genetic basis of mendelian phenotypes: Discoveries, challenges, and opportunities. The American Journal of Human Genetics 97, 199–215 (2015).

30. Badonyi, M. & Marsh, J. A. Prevalence of loss-of-function, gain-of-function and dominant-negative mechanisms across genetic disease phenotypes. bioRxiv 2025–03 (2025).

31. Gerasimavicius, L., Livesey, B. J. & Marsh, J. A. Loss-of-function, gain-of-function and dominant-negative mutations have profoundly different effects on protein structure. Nature communications 13, 3895 (2022).

32. Livesey, B. J. & Marsh, J. A. Why variant effect predictors and multiplexed assays agree and disagree. bioRxiv 2025–07 (2025).

33. Fortuno, C. et al. Specifications of the ACMG/AMP variant interpretation guidelines for germline TP53 variants. Human mutation 42, 223–236 (2021).

34. Parsons, M. T. et al. Evidence-based recommendations for gene-specific ACMG/AMP variant classification from the ClinGen ENIGMA BRCA1 and BRCA2 variant curation expert panel. The American Journal of Human Genetics 111, 2044–2058 (2024).

35. Grimm, D. G. et al. The evaluation of tools used to predict the impact of missense variants is hindered by two types of circularity. Human mutation 36, 513–523 (2015).

36. Harrison, S. M., Biesecker, L. G. & Rehm, H. L. Overview of specifications to the ACMG/AMP variant interpretation guidelines. Current protocols in human genetics 103, e93 (2019).

37. Pathak, A. K. et al. Pervasive ancestry bias in variant effect predictors. bioRxiv 2024–05 (2024).

38. Xu, H. et al. Single cell sequencing as a general variant interpretation assay. bioRxiv 2023–12 (2023).

39. Rowlands, C. F. et al. Availability of benign missense variant truthsets for validation of functional assays: Current status and a novel systematic approach. medRxiv 2025–03 (2025).

40. Axakova, A. et al. Landscapes of missense variant impact for human superoxide dismutase 1. bioRxiv (2025).

41. Findlay, G. M. et al. Accurate classification of BRCA1 variants with saturation genome editing. Nature 562, 217–222 (2018).

42. Zheng, H. et al. Proactive functional classification of all possible missense single-nucleotide variants in KCNQ4. Genome Research 32, 1573–1584 (2022).

43. Jia, X. et al. Massively parallel functional testing of MSH2 missense variants conferring lynch syndrome risk. The American Journal of Human Genetics 108, 163–175 (2021).

44. Lo, R. S. et al. The functional impact of 1,570 individual amino acid substitutions in human OTC. The American Journal of Human Genetics 110, 863–879 (2023).

45. Clausen, L. et al. A mutational atlas for parkin proteostasis. Nature Communications 15, 1541 (2024).

46. Kotler, E. et al. A systematic p53 mutation library links differential functional impact to cancer mutation pattern and evolutionary conservation. Molecular cell 71, 178–190 (2018).

